## Supplemental information for "The histone chaperone NASP maintains H3-H4 reservoirs in the early Drosophila embryo"

### A.

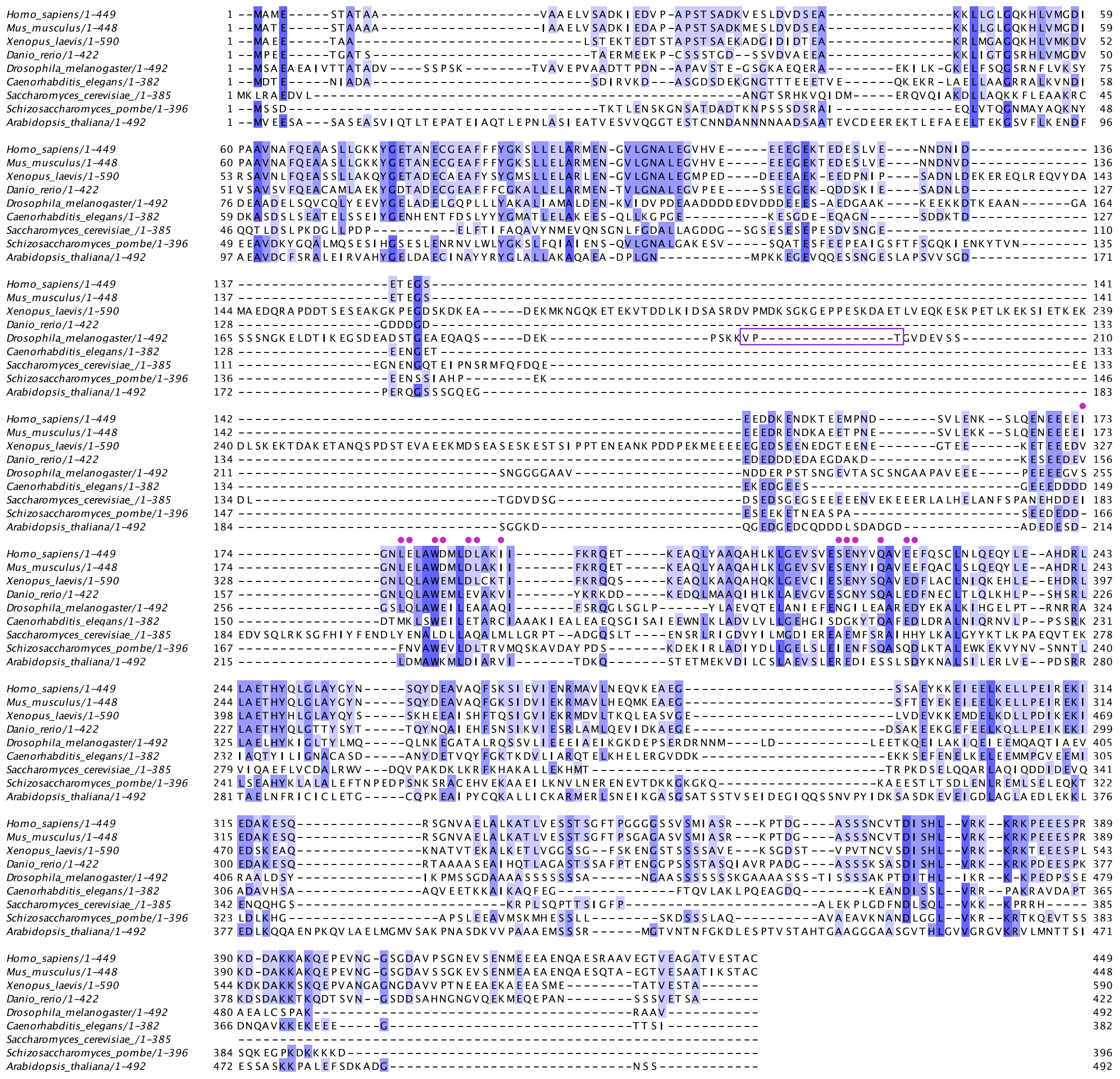

B.

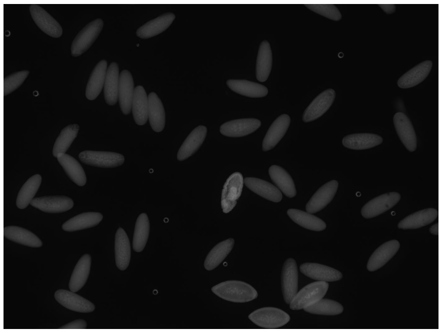

| Replicate | Raw Data | % |
|---|---|---|
| 1 | 47/50 | 94 |
| 2 | 48/50 | 96 |

C.

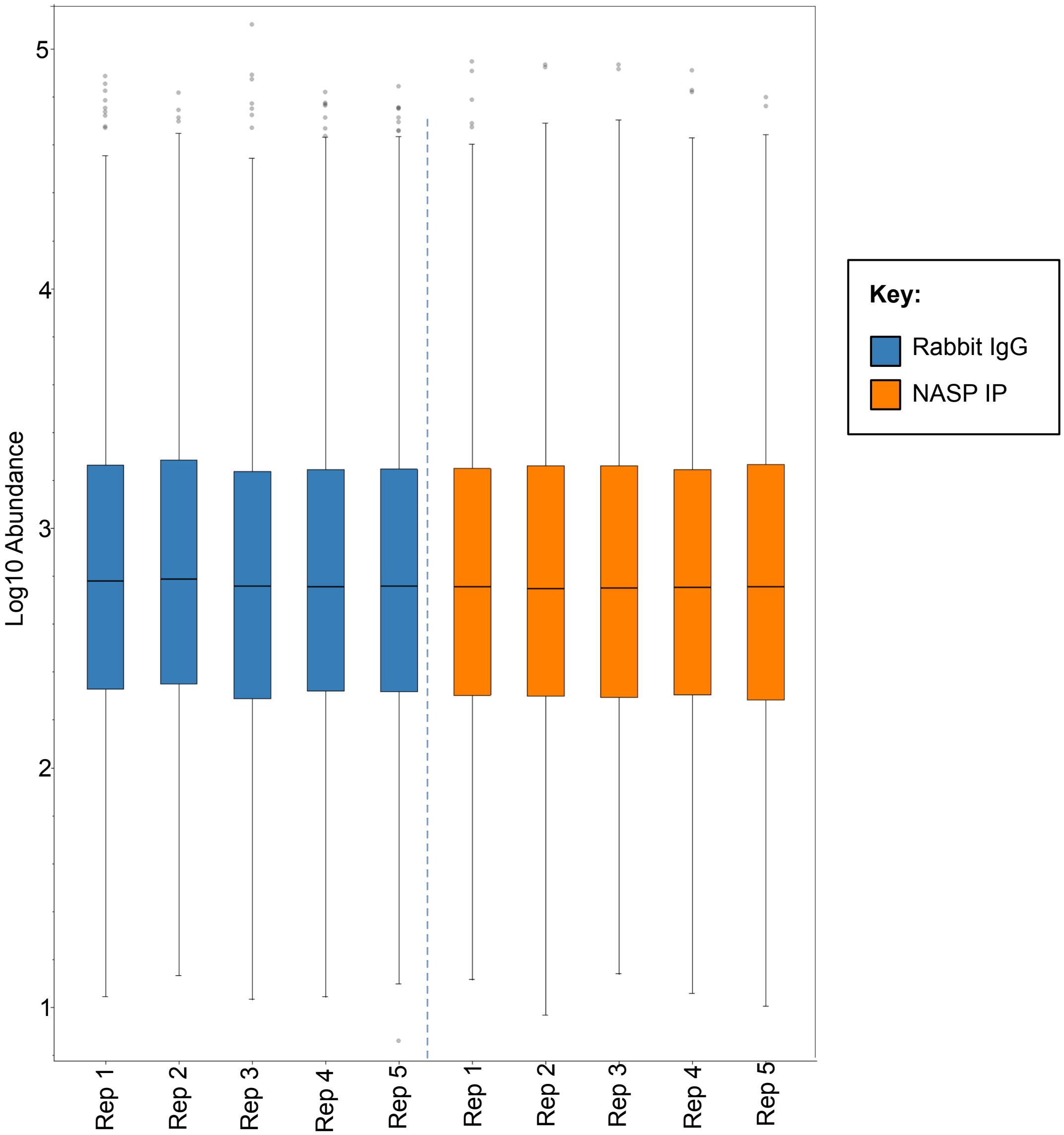

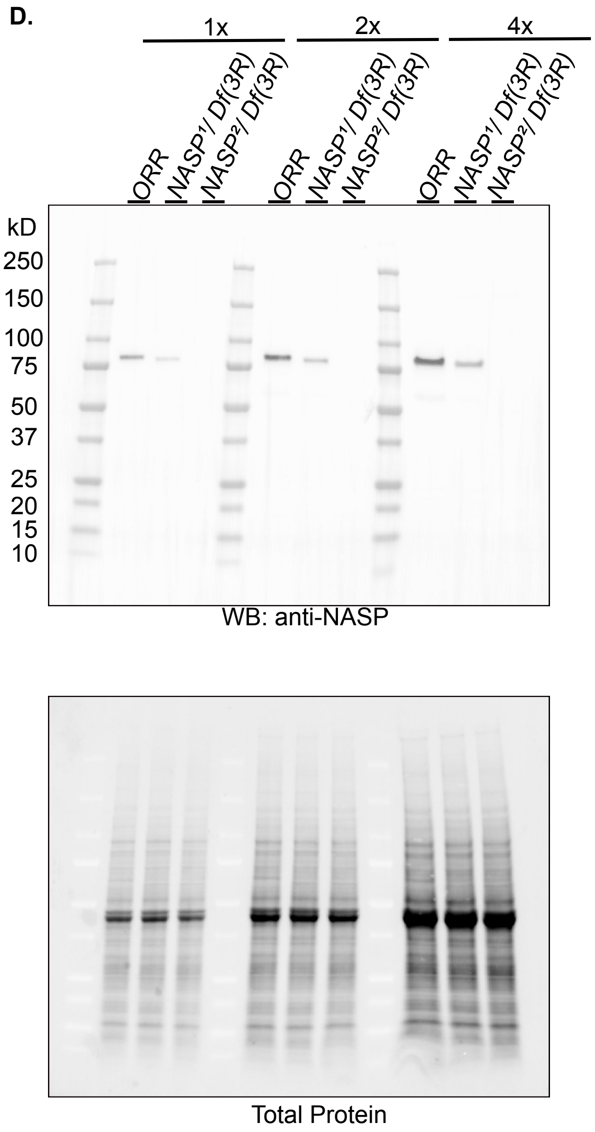


**S1 Fig. Source data for Fig 1**

**(a)** Sequence alignment of CG8223 with NASP homologs in other organisms. Darkening of the color indicates greater conservation. Magenta dots represent the α-N Histone H3 binding region observed in *Homo sapiens* NASP. Boxed region represents the gRNA target sequence for CRISPR-based mutagenesis to generate *NASP* mutants. **(b)** Representative image of DAPI stained 0-2hr AEL embryos with the percentage of stages 1-14 embryos from two replicates. **(c)** TMT abundance of each channel for mass spectrometry IP TMT normalized to total peptides. **(d)** Western blot analysis of ovary extracts from the indicated genotypes with total protein loading control. Cropped blot is used from this full blot for Fig 2B.

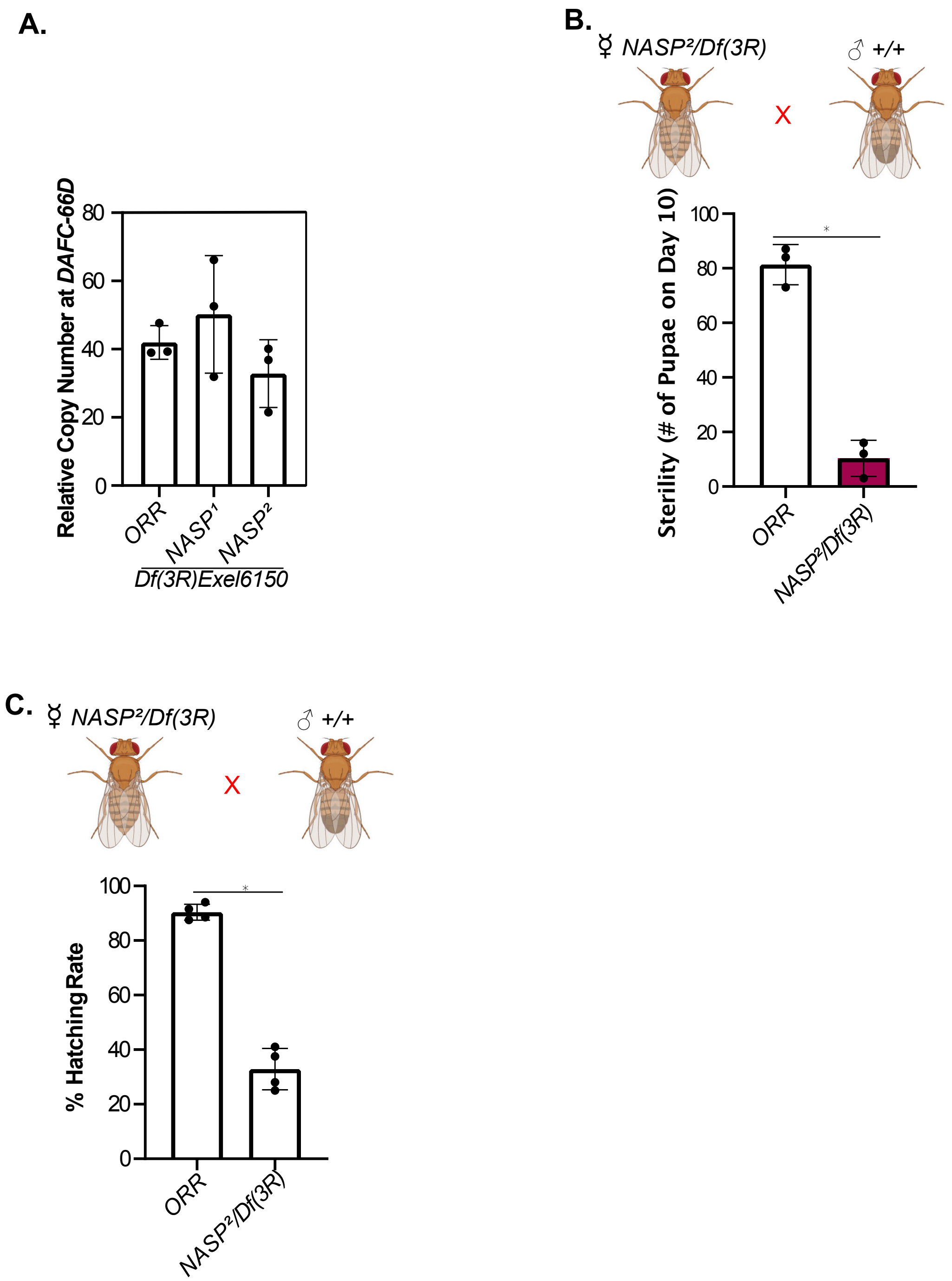


**S2 Fig. Source data for Fig 2**

**(a)** *DAFC-66D* copy number relative to a non-amplified control locus from stage 12 egg chambers for the genotypes listed on the x-axis. Kruskal-Wallis ANOVA was performed to determine significance ($p<0.05$). **(b)** The number of pupae on day 10 produced from virgin females with the genotypes outlined on the x-axis crossed with wild type males. Each data point is representative of a biological replicate ($n=3$). Unpaired t-test was used to determine significance ($p<0.05$). **(c)** Percentage of embryos hatched laid by wild type or *NASP²/Df(3R)Exel6150* mothers. Each data point is representative of a biological replicate ($n=4$) and represents the hatch rate of a group of

100 embryos. Dunn’s Multiple Comparison post-hoc was performed to determine significance ( $p<0.05$).

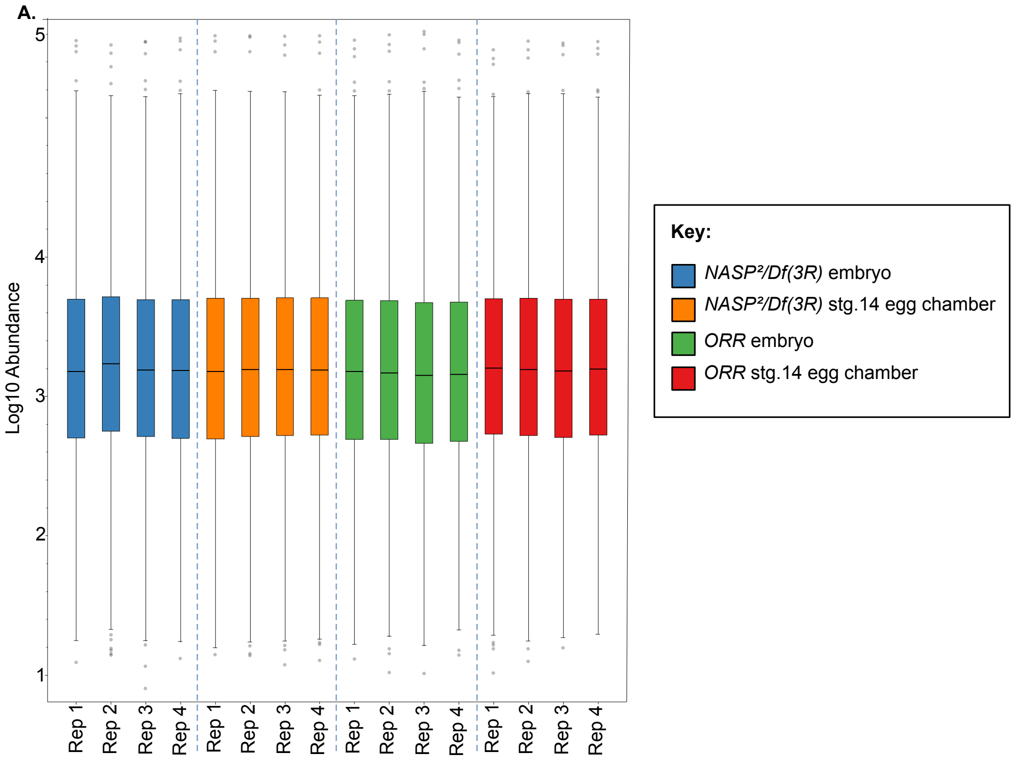


**S3 Fig. Source data for Fig 3**

(a) TMT abundance of each channel for mass spectrometry normalized to total peptides of wildtype or NASP mutant 0-2hr embryos and stage 14 egg chambers.
